# Self-supervised learning yields representational signatures of category-selective cortex

**DOI:** 10.64898/2026.02.09.704031

**Authors:** Daniel Janini, Radoslaw Cichy

## Abstract

The ventral visual stream contains category-selective regions with distinct feature tuning, most prominently the fusiform face area (FFA) and parahippocampal place area (PPA). Why do these brain regions exhibit distinct tuning properties? Recent work suggests that brain-like category-selective features emerge from a general visual learning mechanism without domain-specific biases. Here, we test this proposal by applying a functional localizer approach to both humans and self-supervised neural networks, identifying face- and scene-selective units in the brain and in models. We then compared fMRI and model responses across a broad stimulus set probing classic representational signatures of the FFA and PPA, including preferences relating to curvature, animacy, real-world size, mid-level features, face shapes, and spatial layout information. Category-selective model units largely recapitulate the distinct representational signatures of category-selective brain regions, capturing most of the effects in our test battery. Our findings demonstrate that domain-general learning objectives are sufficient to create humanlike category-selectivity, suggesting that the distinct representational signatures of category-selective cortex may emerge from a unified computational goal akin to self-supervised learning.

## 2. INTRODUCTION

Category selectivity is a hallmark of the ventral visual stream, with separate cortical regions selectively responding to faces^1–3^, scenes^4–6^, bodies^7,8^, and words^9,10^. In particular, the fusiform face area (FFA) and parahippocampal place area (PPA) have been the focus of extensive research, revealing a suite of distinct representational signatures in each region^11–14^. The FFA is driven by the shape of a face, with preferential responses generalizing across a variety of surface properties including animal faces^15,16^, cartoon faces^15^, and illusory faces in objects^17,18^. In contrast, the PPA is primarily driven by the spatial layout of a scene rather than objects^4,19^. Beyond these face- and scene-specific properties, these regions also exhibit distinct preferences related to curvature^20,21^, animacy^22,23^, real-world size^22,24^, and mid-level features^25^. Together these effects constitute a classic set of representational signatures for face- and scene-selective cortex. What drives the emergence of these distinct representational signatures?

One proposal is that representations in the FFA and PPA are optimized for distinct, domain-specific objectives^26,27^. For example, the FFA may be optimized for person identification^28–30^, while the PPA may be optimized for navigational behaviors^31–33^. Consistent with this view, the FFA and PPA are situated within separate brain networks^34–37^. Face-selective cortex is connected to a social processing network including amygdala and medial prefrontal cortex^35,38,39^, while scene-selective cortex is connected to a navigation-related network including hippocampus^40,41^. Some researchers propose that such domain-specific connectivity is the primary cause of category-selective feature tuning^26,27^. In terms of computational evidence, specialized neural networks trained on face categorization align with the perceptual similarity of faces better than general purpose networks trained on object categorization^42,43^. Together, these findings suggest that domain-specific architectural constraints and specialized learning pressures may be required to produce the representational signatures of the FFA and PPA.

An alternative proposal holds that category-selective features emerge from domain-general learning mechanisms^44–46^. Recently, self-supervised neural networks provide a computational model for this idea: they learn image representations tolerant to low-level transformations without labelled feedback or domain-specific architectural biases. Supporting this proposal, self-supervised model features predict neural responses throughout occipitotemporal cortex^47,48^. Regarding category-selectivity, face- and scene-selective features naturally emerge in these networks ^49–51^. Moreover, linear mappings from these category-selective model units to their corresponding brain regions accurately predict neural responses to natural scenes^49,51^. However, it is unknown whether these category-selective model features exhibit the classic representational signatures of the FFA and PPA. Specifically, it is untested whether such models account for sensitivities to curvature, animacy, real-world size, mid-level features, face shapes, and scene layout information. Demonstrating these effects would provide strong evidence for representational alignment between self-supervised models and category-selective brain regions, while the absence of these effects would challenge the plausibility of domain-general accounts of feature learning.

In this study, we tested whether hallmark representational signatures of the FFA and PPA emerge from self-supervised learning. We directly compared feature tuning in models and the brain with a functional localizer approach, identifying the FFA and PPA in fMRI participant, and then applying an analogous procedure to identify face- and scene-selective features in self-supervised networks. We then constructed a test battery of images probing a wide set of classic representational signatures of the FFA and PPA, including sensitivity to curvature, animacy, real-world size, mid-level features, face shapes, and scene spatial layout. Category-selective model features exhibited substantial correlations with their corresponding brain regions and demonstrated the majority of the representational signatures in the test battery. These results indicate that self-supervised model features are aligned with category-selective brain regions, providing computational support for the proposal that humanlike category-selective representations emerge from domain-general learning processes.

## 3. RESULTS

### 3.1 Self-supervised models contain face- and scene-selective units

Our first goal was to identify face- and scene-selective units in self-supervised models which could be compared to the FFA and PPA in subsequent analyses. In fMRI research, face-and scene-selective neural populations are typically identified using a localizer procedure that contrasts voxel responses to faces, scenes, and objects^2,4,52^. Recently, analogous procedures have revealed category-selective representations in category-supervised^42,53–55^ and self-supervised neural networks^49–51^. In the brain, the FFA and PPA have been identified with a wide variety of stimulus sets^2–4,11,22,56^, indicating that their selectivity is not dependent on any single probe set of images. Therefore, we aimed to identify model units which similarly exhibit robust category selectivity across standard localizer image sets. Identifying such units ensures that subsequent model-brain comparisons rely on units with stable category-selectivity rather than units whose preferences are dependent on a particular image set.

We identified category-selective model units in a two-step procedure. First, we identified candidate face- and scene-selective model units in eight self-supervised ResNet50 models trained on ImageNet-1K^57^ (See Methods: Models). We measured unit-wise responses to faces, scenes, and objects in four ReLU layers corresponding to the output of each hierarchical model stage (see Figure 1A for example images). In each layer of each model, we used a conjunction test to identify candidate category-selective units, selecting units with larger responses to the target category than both the other categories (e.g., faces > objects *and* faces > scenes, each p < 10^-3^, t-tests across image activations). Because the selection criteria include a direct contrast between faces and scenes, the resulting sets of face- and scene-selective units are mutually exclusive.

**Figure 1.**
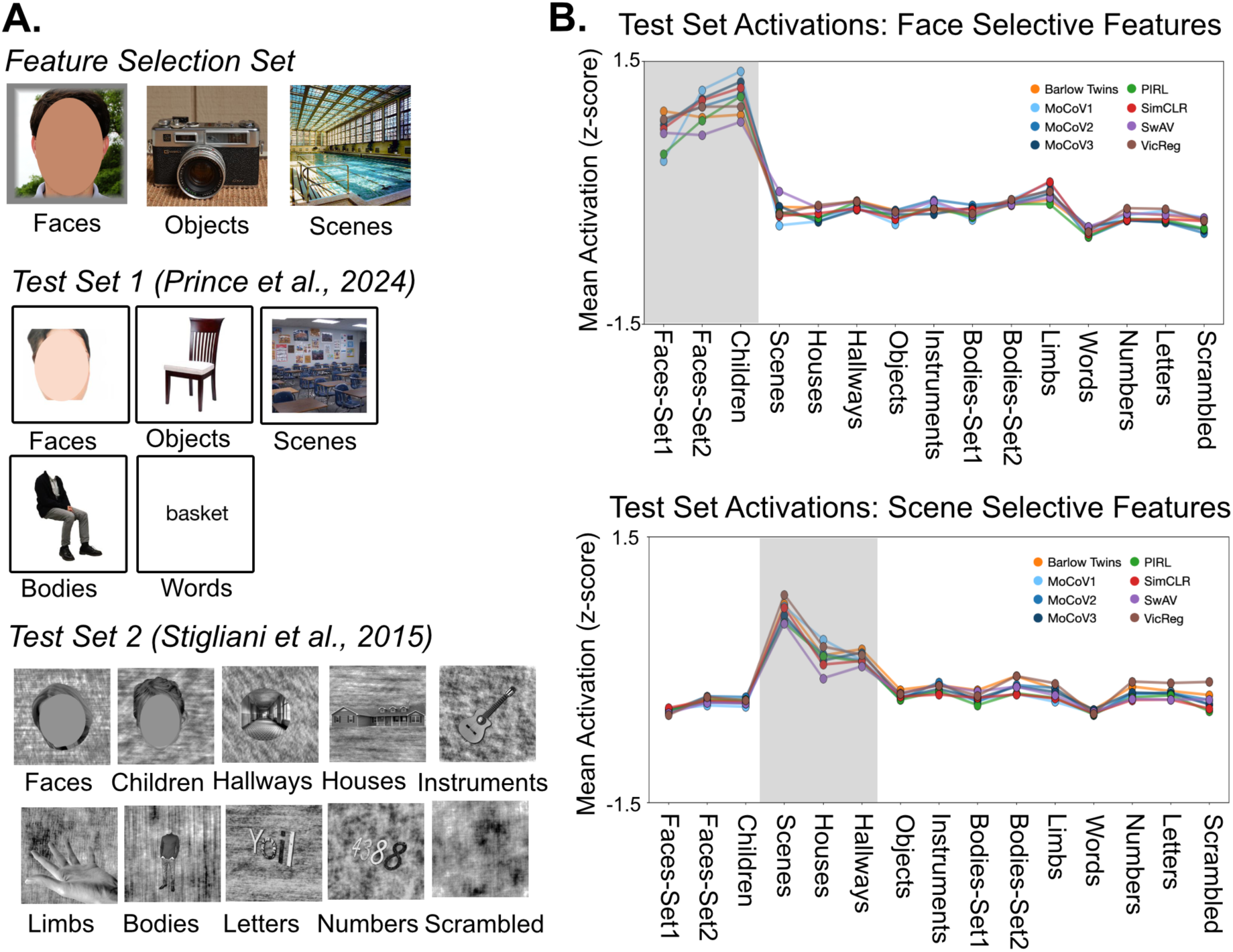
Selecting face- and scene-selective units in self-supervised models. **A**) Example images from the selection set and test sets from the prior literature. The feature selection set was used to identify candidate category-selective units, with scene-selective units responding more to scenes than both faces and objects, and face-selective units responding more to faces than both scenes and objects. Responses to the test sets were used to determine if the units exhibited robust selectivity across standard localizer image sets. Identifiable faces were removed from this figure. **B**) Activations to the test set conditions in the final stage of each network (the ReLU output of Layer 4.2). Each point represents the mean activation across the model units. Images from the target categories are highlighted in the translucent gray area (adult faces and children’s faces for face units; scenes, houses, and hallways for scene units). Face-selective units in every model responded more to face categories than all other categories (p < 10-4 for each pair of conditions, t-tests across image activations). Likewise, scene-selective units in every model responded more to scene categories than all other categories (p < 10-4 for each pair of conditions, t-tests across image activations).

Second, we tested whether these candidate units maintained selectivity for their preferred category across standard fMRI localizer image sets (see Fig. 1A for example images), and we found robust category-selectivity in higher level feature spaces. We compiled two test sets from prior fMRI and modeling studies which vary in low-level properties: 1) a color image set on a white background^49^, and 2) a black and white image set with phase scrambled backgrounds controlling for surface area and luminance properties^49,51,56,58–61^. In the final stage of every model, face- and scene-selective units maintained their selectivity across the test sets (Fig. 1B). The mean response over face-selective units was higher for faces than all other conditions of objects, bodies, scenes, and symbols (p < 10^-4^ for each pair of conditions, t-tests across image activations). Likewise, the mean response over scene-selective units was higher for scenes, hallways, and houses than all other conditions (p < 10^-4^ for each pair of conditions, t-tests across image activations). This establishes that units were consistently category-selective across different image sets varying in low-level properties. Across the networks, 3.6%±0.07% (mean±sd) of the units in this final ReLU layer were face-selective while 3.2%±1.0% of the units were scene-selective.

Consistent with previous research arguing for weaker category preferences in earlier model layers^49^, we found that candidate category-selective units in earlier ReLU layers did not maintain selectivity for their target category across the test sets (Supplementary Fig. 1). Specifically, candidate face-selective units did not selectively exhibit higher responses for faces over all other categories. Likewise, candidate scene-selective units did not selectively exhibit higher responses for scene categories. Thus, in early model layers, units selected for their category preferences in one image set did not maintain selectivity for that category in the test sets.

Next, we applied the same procedure to an untrained ResNet-50 architecture to determine whether architectural constraints alone yield category-selective feature tuning. In contrast to trained networks, previous research has yielded inconsistent results on whether category-selective features exist in untrained networks^49,61^. We identified candidate face- and scene-selective units from the selection image set; however, this selectivity did not generalize in the test sets, indicating a lack of robustly category-selective features in untrained networks (Supplementary Fig. 2). Self-supervised training, rather than just architectural constraints, caused the emergence of robust category-selective features.

Thus, we identified units with robust category-selectivity in the final layer of eight self-supervised models. These results highlight the importance of testing selectivity across multiple stimulus sets when identifying category-selective units in neural network models. Doing so prevents misidentifying units with inconsistent category preferences, for example, in low-level feature spaces or untrained models. Next, we directly compare feature tuning in category-selective model units and category-selective brain regions.

### 3.2 Constructing a test battery for category-selective units and measuring cortical responses

To test whether category-selective model features exhibit major representational signatures of category-selective cortex, we constructed an image set inspired by influential papers on the FFA and PPA. This image set includes 40 hypothesis-driven conditions (5152 images) selected to isolate specific feature preferences. Example images from each condition can be viewed in Fig. 2A. These conditions target effects related to curvature, mid-level features, animacy, real-world size, visual size, face shape tuning, and spatial properties of scenes. The test battery combines the benefits of targeted experimental contrasts with broader sampling across a variety of feature dimensions.

**Figure 2.**
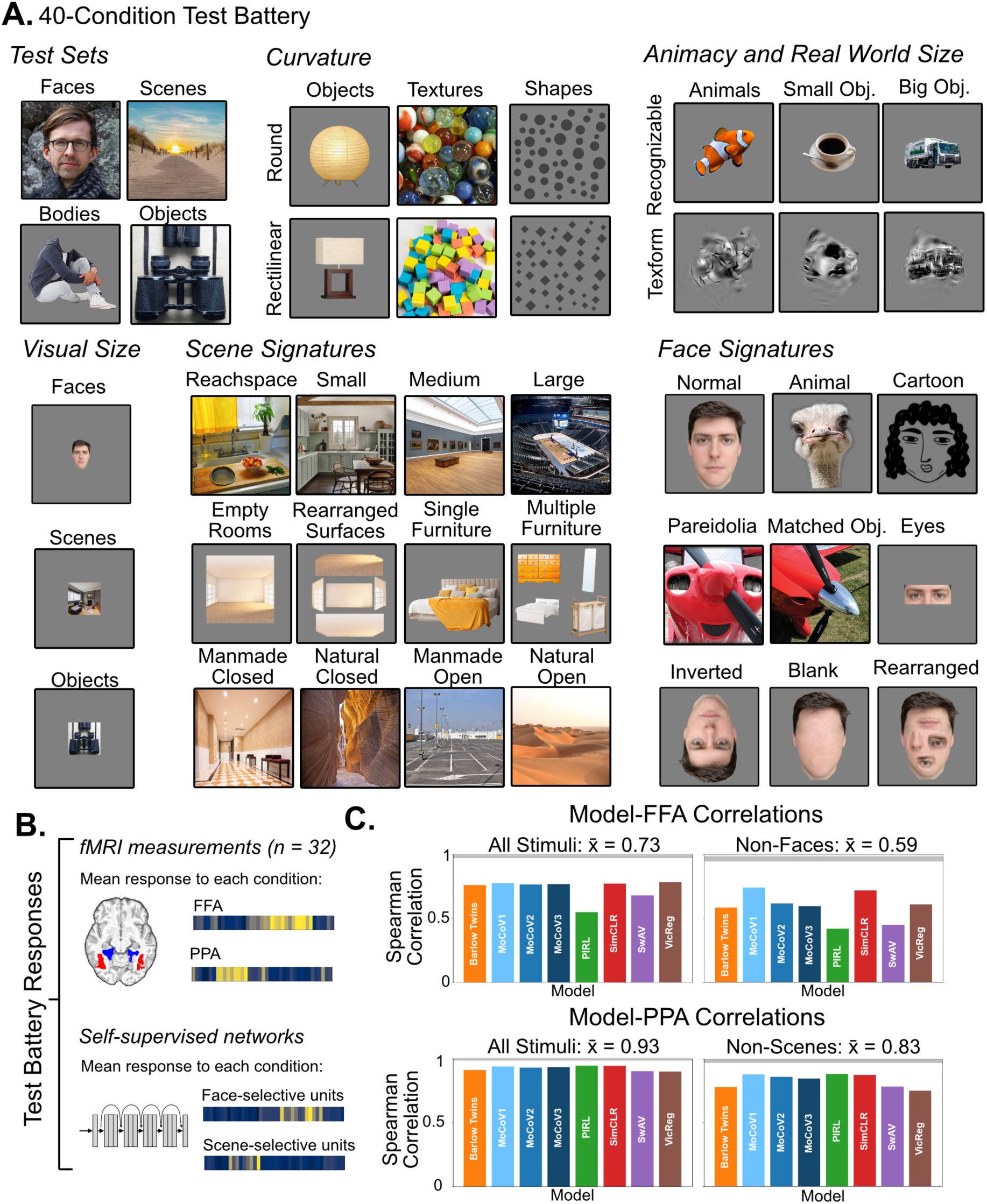
A test battery for face- and scene-selective features. **A**) Example images from each of the 40 conditions in the test battery. Conditions sample a variety of signatures of the FFA and PPA including preferences relating to curvature, animacy and real-world size, visual size tolerance, spatial layout information in scenes, and face shape tuning. Face images included in this figure are of the two authors (RC and DJ). **B**) Responses to the test battery were measured with fMRI and in self-supervised networks. Across 32 participants, the mean response to each condition was measured in the FFA and in the PPA. Likewise, in each self-supervised neural network, the mean response to each condition was measured in the face-selective units and in the scene-selective units. **C**) Correlations were measured between category-selective brain regions and category-selective model units. The gray bar at the top of each plot indicates the noise-ceiling of the fMRI response profiles. Correlations were measured with all conditions, and when excluding the preferred category (e.g., excluding scenes for scene-selective features).

Our central empirical question is whether category-selective model features and category-selective brain regions exhibit similar preferences across this test battery. Addressing this question requires reliable neural measurements to the test battery. Therefore, we collected fMRI responses to each condition of the test battery using a block-design paradigm (n = 32 participants). We defined the FFA and PPA in each participant using the same images and contrasts that we used to identify face- and scene-selective model units. Then we measured the mean activity of the FFA and PPA to each of the 40 conditions (i.e., their response profiles). The response profile measures the graded univariate response of each region across the 40-condition test battery. In both regions, the reliability of the response profile was very high (FFA: ρ = 0.985, PPA: ρ = 0.992; Spearman-Brown corrected reliability across participants). Thus, we obtained highly reliable neural measurements across a rich test battery, providing a stringent upper bound for tests of model-brain correspondence.

### 3.3 Category-selective model features align with category-selective brain regions

We next measured the overall correspondence between category-selective model units and the category-selective brain regions across the full test battery. We measured the mean response to each condition in the face-selective units and in the scene-selective units. This provided a measurement of the graded univariate responses of each set of features across all the test battery condition. We then measured the correlations between these model response profiles and the response profiles of the FFA and PPA. Across the different networks, model response profiles exhibited high correlations with the FFA (mean correlation: ρ = 0.73±0.08, mean±sd) and the PPA (ρ = 0.93±0.02, Fig. 2B). While model-brain correlations were high for both regions, they were higher for the scene-selective units than the face-selective units (p = 0.008, two-sided binomial test). Overall, face- and scene-selective units in the self-supervised models exhibit substantial correspondence to face- and scene-selective regions of the brain.

To test whether model features simply capture a binary distinction between the preferred category and all other categories, we measured model-brain correlations after excluding images for the preferred category. For example, measuring the correlation between face-selective model units and the FFA when only considering images of objects and scenes. Model-brain correlations did decrease after removing the preferred category (p < 10^-4^, two-sided binomial test); however, the correlations were still largely preserved in both the face-selective units (mean correlation with FFA: ρ = 0.59±0.11) and the scene-selective units (mean correlation with PPA: ρ = 0.83±0.05). Thus, model features capture graded variance among non-preferred categories, not just a binary preference for their preferred category.

Thus, category-selective model units and category-selective brain regions exhibit similar preferences across the test battery, supporting the view that domain-general learning produces humanlike category-selectivity. However, these model-brain correlations do not reveal which specific representational signatures are shared between models and category-selective brain regions. To test for specific signatures, we next compared activations to targeted sets of conditions, determining if model features exhibit a suite of classic effects relating to curvature, animacy, real-world size, mid-level features, visual size, face shape tuning, and spatial properties of scenes.

#### 3.4.1 Testing specific representational signatures

To determine if category-selective model units exhibit the same univariate effects as category-selective brain regions, we applied the following approach: First, we assessed whether classic univariate effects from the literature replicated in our fMRI dataset. Second, we measured the replicated effects in the neural network models. Effects which did not replicate in our fMRI dataset were not tested in the models, because it was not clear what would constitute model-brain alignment. To determine if models consistently exhibited the same effects as the brain, we used a prevalence testing framework^62,63^. For each effect, we performed significance testing within each model separately. At a group level, we statistically tested if the prevalence of significant effects across models exceeded chance. If the models not only consistently exhibited significant effects, but also effects in the same direction as the brain, then we concluded that the effect consistently emerged among the models tested.

#### 3.4.2 Model features exhibit brain-like curvature preferences

In the brain, the FFA and PPA exhibit opposite curvature preferences, with the FFA preferring round stimuli and the PPA preferring rectilinear stimuli^5,20,21,64^. These findings indicate that category-selectivity meaningfully covaries with lower-level contour shape preferences, rather than being fully abstracted from these more primitive visual features. We tested this effect with three types of stimuli from prior work: 1) textures (e.g., bubbles vs metal gratings), 2) isolated objects matched in their real-world size (e.g., round vs rectilinear chairs), 3) two-dimensional arrays of circles and diamonds.

In our fMRI dataset, we replicated curvature effects for both texture and object stimuli (Fig. 3A). The FFA responded more to curvy stimuli than rectilinear stimuli (texture: p = 0.0001, objects: p = 0.0005, two-sided binomial tests). In contrast, the PPA responded more to rectilinear stimuli than curvy stimuli (textures: p < 10^-4^, objects: p= 0.05, two-sided binomial tests). However, neither region exhibited a significant curvature effect for two-dimensional shapes (FFA: p = 0.38, PPA: p = 1.0, two-sided binomial tests). Thus, curvature preferences were observed in the FFA and PPA for naturalistic stimuli, but not for simple geometric shapes.

**Figure 3.**
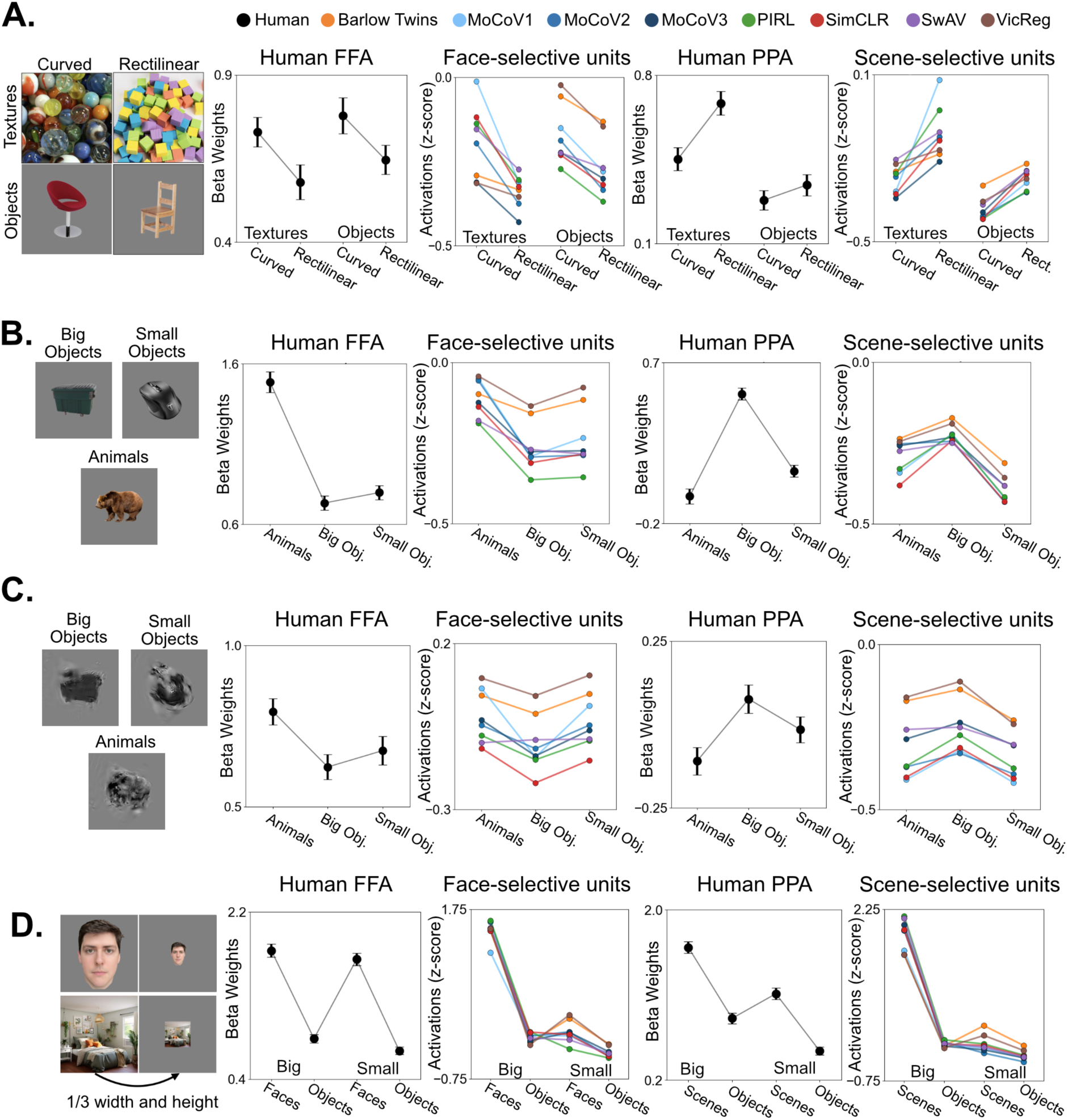
Testing representational signatures of the FFA and PPA. A) Curvature preferences. Left, Example images of curved and rectilinear stimuli. Right, mean activation to each condition in the human FFA and PPA and in category-selective model features. B) Animacy and real-world size preferences. Left, example images of each condition. Right, mean activation to each condition in the human FFA and PPA and in face- and scene-selective model units. C) Mid-level features of animal and object conditions. Left, example texform images, which preserve second-order spatial frequency statistics of the original images, but are unrecognizable. Right, mean activation to each texform condition. D) Visual size versus category effects. Left, example images at two visual sizes. For human participants, image content was displayed at either 2.25 or 6.75 degrees of visual angle. Right, mean activation to categories at different visual sizes. Face images included this figure are of author DJ.

We found similar curvature preferences for textures and objects in self-supervised neural networks (Fig. 3A). In every network, face-selective units responded more to curvy textures than rectilinear textures and more to curvy objects than rectilinear objects (p < 10^-4^, population prevalence testing). Scene-selective units exhibited an opposite preference for rectilinear textures and objects (all p < 10^-4^, paired t-tests in each model). As in the brain, face- and scene-selective units exhibit opposite curvature preferences. Thus, covariation between curvature preferences and category-selectivity naturally emerges from domain-general visual learning processes.

#### 3.4.3 Model features exhibit brain-like animacy and real-world size preferences

The human FFA and PPA are situated within broader cortical topographies related to animacy and real-world size. While most selective for faces, the FFA overlaps a region preferring animals over big and small objects^22,23,65^. Also, even though the PPA is most selective for the spatial structure of a scene rather than objects, it still prefers big objects over small objects and animals^22,24^. These findings indicate that face- and scene-selective neural populations are meaningfully situated within major joints of the representational structure of objects.

We replicated these representational signatures by comparing fMRI responses to animals, big objects, and small objects (Fig. 3B). The FFA exhibited a higher response to animals than big objects (p < 10^-4^, two-sided binomial test) and small objects (p < 10^-4^, two-sided binomial test). The PPA exhibited a higher response to big objects than small objects (p < 10^-4^, two-sided binomial test) and animals (p < 10^-4^, two-sided binomial test). Category-selective model units exhibited similar animacy and real-world size preferences as category-selective brain regions. Face-selective units responded significantly more to animals than big objects (p < 10^-4^, population prevalence testing) and more to animals than small objects (p < 10^-4^, population prevalence testing). Scene-selective model units responded significantly more to big objects than small objects (p < 10^-4^, population prevalence testing), and more to big objects than animals (p < 10^-4^, population prevalence testing).

Thus, self-supervised models capture how face- and scene-selective features covary with animacy and real-world size preferences.

In the human visual system, animacy and real-world size preferences are partially driven by the mid-level features of stimuli^25,66–68^. Prior research indicates that cortical preferences for animacy and real-world size are maintained even when stimuli are converted to “texforms”, which preserve second-order spatial frequency statistics^69^ but obscure higher level features needed to recognize images at a basic category level^25,68^ (see Fig. 3C for examples). We generated texform versions of the animals, big objects, and small objects, and we found that animacy and real-world size preferences were maintained with these stimuli. The FFA responded more to animal texforms than to big object texforms (p < 10^-4^, two-sided binomial test) and more to animal texforms than to small object texforms (p = 0.02, two-sided binomial test). The PPA responded more to big object texforms than to small object texforms (p = 0.02, two-sided binomial test) and more to big object texforms than to animal texforms (p = 0.002, two-sided binomial test).

Next, we tested texform responses in the self-supervised models to determine if animacy and real-world size preferences are similarly maintained by second-order spatial frequency statistics. Face-selective units responded significantly more to animal texforms than big object texforms (p < 10^-4^, population prevalence testing), and more to animal texforms than small object texforms (p = 0.0004, population prevalence testing). Scene-selective units responded more to big object texforms than small object texforms (p < 10^-4^, population prevalence testing), and more to big object texforms than animal texforms (p < 10^-4^, population prevalence testing).

Thus, category-selective model units are not only embedded within broader representational axes related to animacy and real-world size, but the same mid-level visual features contribute to this organization in both self-supervised models and the human brain. Along with the previous results on curvature preferences, these results indicate that category-selectivity is integrated into a broader mid-level feature space, and that this organization emerges from self-supervised learning mechanisms.

#### 3.4.4 Model features exhibit partial visual size tolerance

One of the primary functions of the ventral stream is to yield visual representations which are tolerant to low-level variation^70,71^. For example, experiments using fMRI and neurophysiological techniques indicate that category-selective regions maintain their selectivity under substantial variation in the visual size of stimuli^72–74^. To replicate these findings, we recorded responses to faces, scenes, and objects at two visual sizes. In one set of images, the content filled the full image surface area similarly to the selection images used to localize the FFA and PPA. In another set of images, the width and height of the image content was 1/3 of the original size and the rest of the image was a uniform gray (see Fig. 3D for example images). For fMRI participants, the image content was presented with corresponding proportions at either 6.75 or 2.25 degrees of visual angle.

We found substantial visual size tolerance in category-selective brain regions (Fig. 3D). For the FFA, responses were higher to faces than objects at both the large visual size and the small visual size (both p<10^-4^, two-sided binomial tests). Likewise, for the PPA, responses were higher to scenes than objects at both visual sizes (both p<10^-4^, two-sided binomial tests). To compare the relative effects of visual size and category, we fit a general linear model to the region of interest condition activations, with categorical variables for visual size and category. In this linear model, the beta weight for category was significantly larger than the beta weight for visual size (p < 10^-4^, t-test comparing beta weights from the linear model). Thus, in the FFA and the PPA, category-preferences outweighed a 9-fold change in image surface area.

To exhibit the same properties as category-selective cortex, model units should maintain their preferences at each visual size, and the effect of category should be greater than the effect of visual size. Face-selective units responded more to faces than objects at each visual size (p < 10^-4^, population prevalence testing). Likewise, scene-selective units responded more to scenes than objects at each visual size (p < 10^-4^, population prevalence testing). However, the networks did not consistently show a greater effect of category than size. For face-selective units, four networks did exhibit a greater category effect, one network exhibited a greater size effect, and in three networks these effects did not significantly differ (threshold of p < 0.05 in each model, t-tests comparing beta weights of linear model). For scene-selective units, two networks exhibited a greater category effect, but five networks exhibited a greater size effect (p < 0.05, t-tests comparing beta weights of linear model).

Thus, self-supervised learning produced features which maintain their preferences at each visual size, but category effects did not consistently outweigh visual size effects like in category-selective brain regions.

#### 3.4.5 Scene-selective model units exhibit partial sensitivity to spatial layout

Next, we assessed a series of effects specific to the tuning of the PPA. The selectivity of the PPA is driven by visual information on the spatial layout of the local environment^4,19^. We replicated three effects in support of this view in our fMRI dataset (Figure 4A). First, the PPA responded more to images of empty rooms than images of the same surfaces rearranged into an incoherent configuration (p < 10^-4^, two-sided binomial test). Second, the PPA responded more to empty rooms than images with single pieces of furniture (p < 10^-4^, two-sided binomial test). Third, the PPA responded more to empty rooms than multiple pieces of furniture (p < 10^-4^, two-sided binomial test). These findings show that the features of the PPA are most responsive to coherent global layout information rather than local surface features or object content.

**Figure 4.**
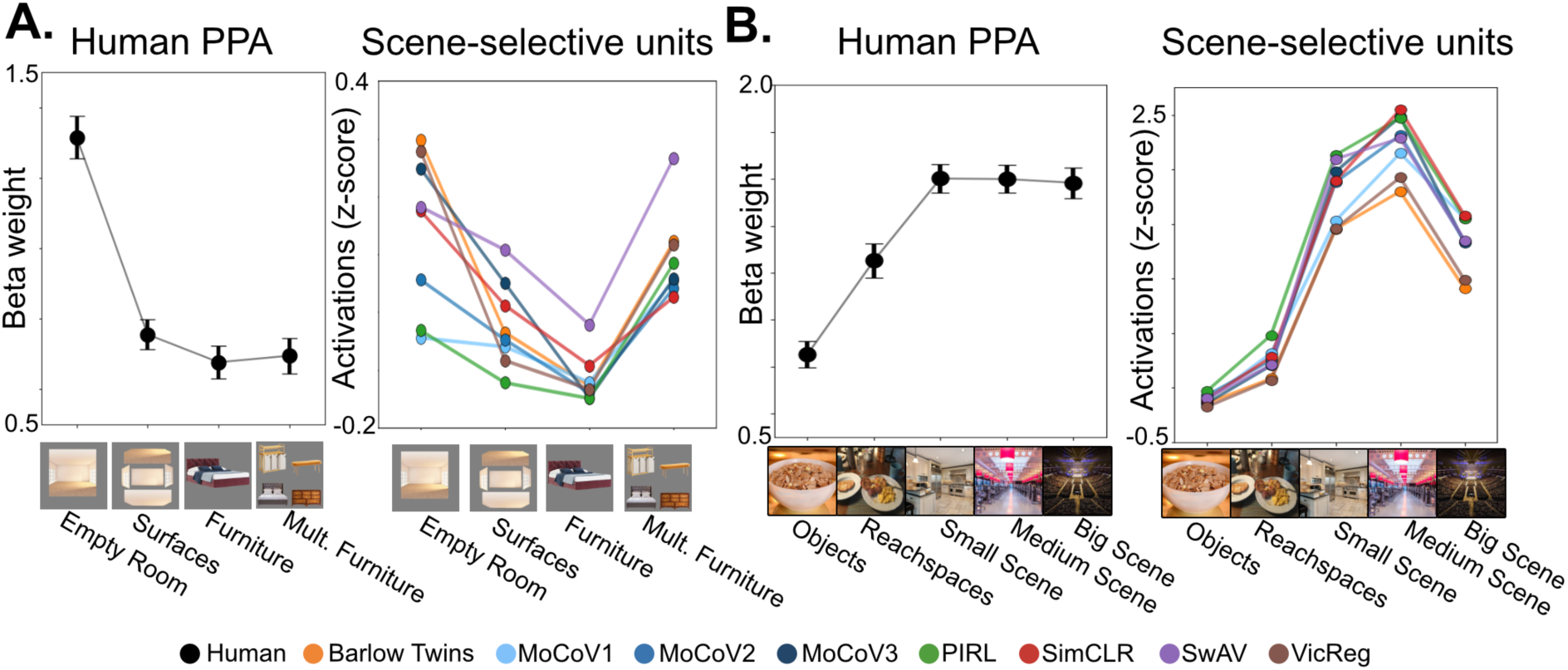
Testing scene-specific signatures. A. Tests for sensitivity to spatial layout information in the human PPA and in scene-selective model units. Responses were measured to empty rooms, room surfaces, furniture, and multiple pieces of furniture. The human PPA responds more to empty rooms than each of the other categories. Scene-selective units responded more to empty rooms than surfaces or single pieces of furniture, but not multiple pieces of furniture. B. Tests for spatial scale in the human PPA and in scene-selective model units. Both the human PPA and scene-selective units responded more to reachspaces than objects, and more to navigable scenes than reachspaces.

Responses in scene-selective model units exhibited 2/3 of these effects. Scene-selective model units responded more to empty rooms than their rearranged surfaces (p < 10^-4^, population prevalence testing) and single pieces of furniture (p < 10^-4^, population prevalence testing). Scene-selective model units consistently exhibited different magnitudes of response to empty rooms and multiple pieces of furniture (p < 10^-4^, population prevalence testing), but the direction of this effect was not consistent: 3/8 networks responded significantly more to empty rooms (p < 0.05, paired t-tests), but 4/8 significantly responded more to multiple pieces of furniture (p < 0.05, paired t-tests), and one network showed no significant difference.

This pattern of results demonstrates that scene-selective model units are not simply responding to local surface properties or singular large objects, indicating a degree of tuning for coherent spatial layout information. However, in comparison to scene-selective cortex, the model units are more biased toward the object content of scenes.

Next, we investigated prior findings claiming that the PPA prefers views of larger scales of space rather than views of nearby space^75–77^. Across prior studies, researchers have measured responses ranging from single object views, to views of reachable surfaces (e.g., a desk), to views of increasingly larger scenes (e.g., a bathroom versus a concert hall). This prior work indicates that responses in the PPA increase across this full range of spatial scales^75–77^. Here, we measured responses in the PPA to objects, reachable surfaces, small scenes on the scale of a bathroom, medium scenes on the scale of an art gallery, and large scenes on the scale of a concert hall (Fig 4A). We successfully replicated the finding that the PPA responded more to reachable surfaces than objects (p < 10^-4^, two-tailed binomial test) and more to navigable scenes than reachable surfaces (p < 10^-4^, two-tailed binomial test). However, the PPA did not exhibit significant differences in its mean response to navigable scenes of different scales (p > 0.05 for paired comparisons between small, medium, and large scenes, two-tailed binomial tests). Our results affirm that the PPA does prefer larger scales of space and not just any space with multiple objects (like a reachable surface), but we did not replicate the finding that the PPA prefers larger scales of navigable space.

Scene-selective model units mirrored the PPA, responding more to reachable surfaces than objects (p < 10^-4^, population prevalence testing). In addition, model units responded more to small, medium, and big scenes than reachable surfaces (p < 10^-4^, population prevalence testing). Thus, both scene-selective model units and the PPA increase their response from views of objects to views of reachable surfaces to views of navigable scenes. As in the brain, scene-selective model units do not just prefer any space with multiple objects, but specifically images of navigable-scale spaces.

Because we did not replicate previous fMRI findings that the PPA prefers big scenes over small scenes, we did not have strong predictions of what would constitute brain alignment in the scene-selective model units concerning variance among navigable scales of space. However, we note that scene-selective units responded to big scenes less than medium and small scenes (p < 10^-4^, population prevalence testing).

Next, we conducted an exploratory analysis to test whether responses in scene-selective model units reflected the distribution of scene types present in the ImageNet training diet. In a random sample of 1000 images from ImageNet, 34.6% of the images depicted a navigable scene. Only 0.2% of the full sample depicted an empty room. Thus, partial preferences for empty room images emerged despite a relative paucity of these images in the training set. Big scenes were underrepresented (7.7%) relative to medium scenes (10.2%) and small scenes (16.7%).

#### 3.4.6 Face signatures: face shapes and parts

Next, we tested signatures in the FFA related to face shape tuning. Face-selective cortex prefers faces across great texture variation, for example maintaining preferential responses for animal faces^15,16,65^, illusory faces in objects (i.e., pareidolia)^17,78^, and cartoon faces^15^. In addition, eyes alone are sufficient to drive preferential responses in the FFA, indicating a particular role for salient face parts^15^. To test these signatures in the FFA, we determined if responses to animal faces, pareidolia faces, cartoon faces, and eyes were all greater than responses to objects. For these comparisons, the object condition included images which had the same categories as the pareidolia faces, but which did not create an illusory percept of a face. All previous findings replicated (Fig 5A). In comparison to the object condition, the FFA exhibited a greater response to animal faces (p < 10^-4^, two-tailed binomial test), pareidolia faces (p < 10^-4^, two-tailed binomial test), cartoon faces (p < 10^-4^, two-tailed binomial test), and eyes (p < 10^-4^, two-tailed binomial test).

**Figure 5.**
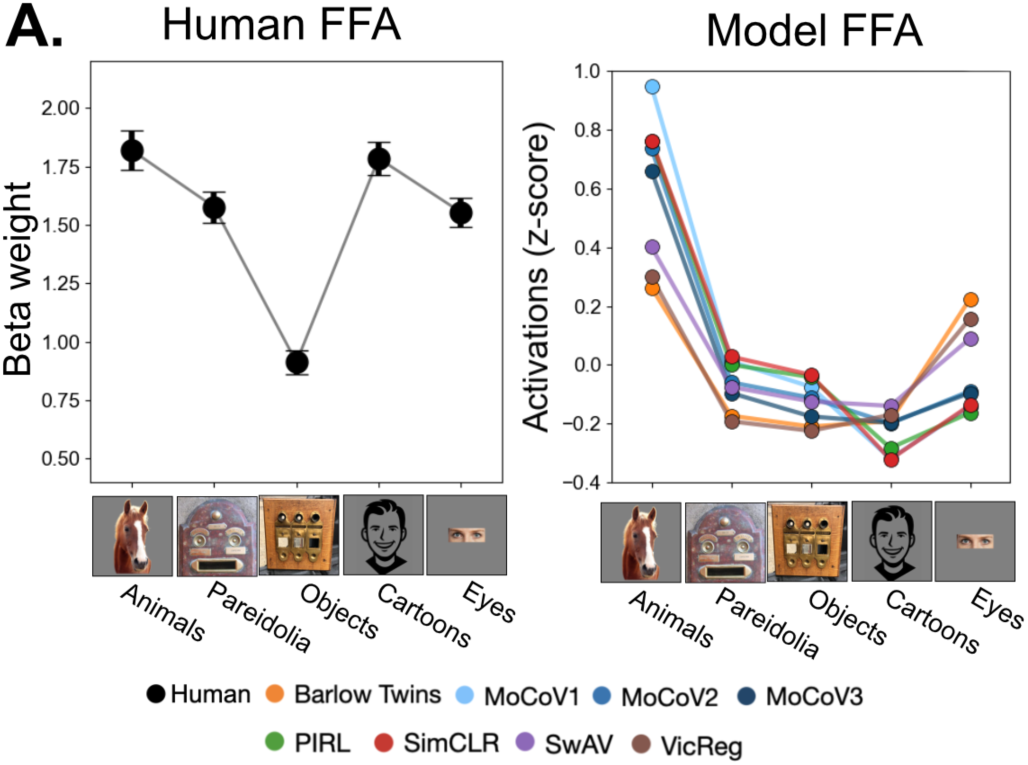
**A**. Face specific conditions measuring sensitivity to face shapes across texture variation. Responses were measured to animal faces, pareidolia faces, objects matched to the pareidolia faces, cartoon faces, and eyes. Left, responses in the human FFA. The FFA responded more to all face conditions than pareidolia-matched objects. Right, responses in face-selective model units. Networks responded more to animal faces and pareidolia faces than pareidolia-matched objects. However, networks did not exhibit consistent preferences for cartoon faces or eyes in comparison to the object condition.

We found that face-selective model units exhibited 2/4 of these effects. Face selective units responded more to animal faces and pareidolia faces than the pareidolia-matched objects (p < 10^-4^ for each comparison, population prevalence testing). However, only one network responded significantly more to cartoon faces than the object condition (p = 0.0008, paired t-test), and four networks instead showed the opposite preference (p < 10^-4^, paired t-test). Networks also differed in their sensitivity to eyes. Four networks responded significantly more to eyes than objects (p<10^-3^, paired t-test), but three networks responded significantly more to objects than eyes (p<10^-3^, paired t-test). The model successes concerning animal faces and pareidolia faces indicate that the features generalize across varied textures rather than being fully texture-biased^79,80^. However, the failure to maintain selectivity for cartoon faces and eyes indicate incomplete face shape tuning.

## 4 DISCUSSION

Longstanding debates concern whether category-selective representations in the ventral visual stream emerge from separate domain-specialized mechanisms or from a general purpose learning process ^26,27,45,81–85^. Recent computational studies indicate that self-supervised learning yields face- and scene-selective visual features^49–51^, but it remains unclear whether these features exhibit classic effects of the FFA and PPA. To address this gap, we constructed a test battery probing canonical effects of the FFA and PPA and directly compared responses between category-selective model units and category-selective brain regions. Across the full test battery, category-selective model units exhibited substantial correlations with their corresponding brain regions and captured most of the tested effects, including preferences related to curvature, animacy, and real-world size. Some effects were partially accounted for including visual size tolerance, the PPA’s sensitivity to spatial layout information, and the FFA’s sensitivity to face shapes. Together, these findings demonstrate that self-supervised learning yields category-selective features largely aligned with the tuning properties of category-selective brain regions. This study provides computational plausibility for domain-general learning accounts of ventral stream representations.

This study demonstrates the power of a functional localizer approach for directly comparing feature tuning in neural networks and the brain. In fMRI research, functional localizers have proven highly useful for identifying common neural populations between participants and characterizing their feature tuning^86^. Here, we show that this same approach provides a simple and powerful framework for model-brain comparison, enabling direct comparisons between category-selective model units and category-selective neural populations. While alternative comparative approaches such as linear encoding models are useful for predicting as much variance as possible in neural data^87^, the functional localizer approach imposes tighter constraints on the mapping between models and the brain. Encoding models typically permit flexible combinations of model features^47,88,89^, including negatively weighted features, and the exclusion of features unaligned with the brain. Researchers have thus called for more constrained and theoretically driven methods for testing representational alignment between model features and the neural responses^47,49,90^. The functional localizer approach directly compares model and brain units with matched selectivity, providing an interpretable and straightforward basis for model-brain comparisons.

We found particularly strong correspondence between scene-selective units in self-supervised networks and the PPA. Prior research on scene perception has often focused on domain-specialized objectives, for example, leveraging networks trained on scene labels and spatial information^91–96^, or theorizing that scene representations are tuned to represent action affordances and navigationally relevant information^11,27,93,97^. Our findings question the need to invoke domain-specific optimization to account for feature tuning in the PPA. However, our models exhibited a notable limitation: an incomplete preference for empty rooms and a bias instead toward arrays of objects. One possible explanation is the object-centric nature of the ImageNet training set, which rarely includes scene surfaces without object content. In contrast, the image diet of a human infant preferentially includes scene views with simple edge statistics such as walls, doors, and ceilings^98^. Future studies could test whether variations in image diet increase preferences for spatial layout information relative to object content.

In our study, face-selective units in self-supervised networks exhibited most of the tested signatures of the FFA, but the model-brain correlations were not as strong as scene-selective features, raising the question of which learning pressures would improve brain alignment. Prior research indicates that both broad visual experience and face-specific learning signals are important. Self-supervised networks and object-categorizing networks better predict neural responses in face-selective cortex than models trained only on face categorization^46,49,99,100^. However, networks trained on objects alone are poor at face categorization^42^, and training on face categorization aligns network representations to behavioral measurements of face similarity^42,43^. These findings indicate that the ideal model would learn to distinguish between faces while situating those representations in the context of a broader visual diet. Future studies could use our test battery to compare the relative impacts of finetuning on face categorization versus self-supervised learning with a richer image diet of faces.

To clarify our argument, we claim that category-selective features emerge from domain-general learning mechanisms, not that category-selective features lack domain-specific functional roles. We do not challenge evidence that the FFA and PPA are situated within separate cortical networks that serve different behavioral aims^34–41,101^. Instead, we join others in differentiating between the learning processes which create visual representations and their functional roles^49^. In fact, recent modelling work has begun to show that modular network structure can naturally emerge from domain-general learning processes, including clustered topographical regions^51,53,54^ and segregated feature populations useful for different tasks^49^. General visual learning mechanisms can produce a rich bank of visual features with varied representational signatures useful for different functional roles.

Compared to human brains, the networks exhibited insufficient visual size tolerance. How is this to be explained? Because the networks were already trained with extensive cropping augmentations (8-100% of the image), this limitation is unlikely to reflect insufficient variation in spatial scale during training. One possibility is that networks lack humanlike mechanisms which distinguish object representations from background information, like foreground segmentation processes and global shape tuning^79,80,102–105^. Therefore, the uniform gray background of the visually small images may have overly influenced feature activations. Another possibility concerns effective receptive fields. The learned weights of a model can restrict the portion of the image that influences a unit^106^, limiting a unit’s ability to generalize across portions of an image. Convolutional architectures may exacerbate these limitations, and transformer architectures may better support global shape representations^107,108^. Future modeling work could investigate how different learning constraints jointly influence receptive field structure, shape representations, and scale tolerant category-selectivity.

Our test battery and modelling framework provide a foundation for investigating other topics relating to category-selectivity. While we focused on faces and scenes due to their well characterized representational signatures, an emerging body of work leverages the localizer approach to investigate model representations of objects, written symbols, bodies, and tools^109–112^. Another topic concerns the emergence of category-selectivity in models and the brain. In visual cortex, category preferences emerge in the first few months of life^113–115^ and continue to develop throughout childhood^116–119^. Competing accounts emphasize domain-specific learning pressures^28,39^, or an initial developmental period of self-supervised learning^120^. Future neuroimaging studies could use our test battery to track the emergence of category-selective signatures across human development and determine which types of networks exhibit similar learning trajectories.

A limitation of this study is that we did not evaluate how domain-specific learning can shape category-selective representations. As such, our findings demonstrate that domain-general forms of learning are sufficient to produce many signatures of category-selective cortex, but they do not necessitate that this type of learning is uniquely capable of doing so. Competing neural network models could operationalize domain-specialized learning pressures through different factors^121^. One factor is image diet, for example training models exclusively on a specific domain (e.g., faces only) or with enriched exposure to certain categories (e.g., broader sampling of scene categories). Another factor is the learning objective itself, with multiple tasks to consider for each domain. For example, scene-related tasks could include categorization, navigational affordance prediction, or construction of surface geometries. Future studies could also compare purely domain-specialized models with models that combine self-supervised pre-training and domain-specialized finetuning. The test battery introduced in this paper provides shared evaluation criteria for comparing such models, paving the way to broader model comparison.

In conclusion, by applying a functional localizer approach to neural network models, we demonstrate that self-supervised learning yields visual features which recapitulate hallmark signatures of category-selective cortex. We join others in providing computational plausibility for the idea that category-selective neural populations emerge as facets of a broader representational space shaped by general purpose learning mechanisms^49–51^. By sampling a broad array of targeted experimental contrasts, the test battery introduced in this study provides a direct and interpretable framework for evaluating the model-brain alignment of category-selective representations.

## 5 METHODS

### 5.1 Models

We evaluated eight self-supervised convolutional neural networks with a ResNet50 architecture: Barlow Twins^122^, MoCoV1^123^, MoCoV2^123^, MoCoV3^124^, PIRL^125^, SimCLR^126^, SwAV^127^, and VicReg^128^. All models were trained on the ImageNet-1K database^57^ and learned feature hierarchies which minimize representational distances between multiple augmentations of the same image. These models rely on the following augmentations: cropping and resizing, horizontal flipping, color jitter, and grayscale transformations. Some networks directly implement a contrastive loss to maximize distances between different images (SimCLR, MoCoV1-3, and PIRL), while other networks implicitly accomplish this goal by minimizing redundancy between features (Barlow Twins, VicReg) or through cluster assignments (SwAV). As models of a general-purpose learning algorithm, these networks did not receive labelled feedback on any domains of visual input, and their innate architecture is general purpose convolutional feature hierarchy.

### 5.2 Stimulus Sets

#### 5.2.1 Localizer image sets

We created an image set of faces, objects, scenes, and bodies to identify face- and scene-selective units in the models, and face- and scene-selective regions in the brain. Each category included 300 images. Images were evenly split between a selection set and a test set. The selection set was used to select candidate face- and scene-selective units, while the test set became part of the test battery. We sampled face from the FlickrFaces database^129^. To create mild position variance of face features in the stimulus set, we randomly resized faces to 49-100% of the image surface area and placed them in a random position on a uniform gray background. Scene images included a variety of indoor and outdoor scenes of different spatial scales. We sampled object images randomly from the Things database^130^. We only included object images with a viewpoint that lacked a clear scene context. Body images included men and women in various action poses (sitting, running, falling, etc.) placed on a gray background in random positions within the image. We removed heads from each body image.

In addition, we tested networks for robustness on two standard localizer image sets from the literature. The first image set includes faces, scenes, objects, bodies, words (80 images per category)^49^. Images are in color, and faces, objects, and scenes are cropped on white backgrounds. The second image set includes adult faces, children’s faces, houses, hallways, cars, stringed instruments, limbs, number strings, letter strings, and phase scrambled images^56^. There are 144 images per category, all in black and white. Image content has position and size variance, and the background in each image is a phase-scrambled image randomly selected from the whole set.

#### 5.2.2 Test Battery

We created an image set sampling conditions from the fMRI literature on the FFA and PPA. This image set includes 40 categories and 5152 images total (see Figure 2A for example images). Below is a description of each of these conditions.

Four conditions were the test sets of faces, objects, scenes, and bodies from the localizer image set described above (150 images each).

Six conditions related to curvature, following prior work on curvature preferences of category-selective regions^20,21^: rectilinear vs round objects (160 images each), rectilinear vs round textures (160 images each), and rectilinear vs round 2D shapes (100 images each). We sampled object images from 64 categories (e.g., book, desk, lamp, tire). We cropped objects and placed them on a gray background. We equated real-world size between rectilinear objects and round objects. We sampled texture images from 32 categories (e.g., marble, wood grain, sugar cubes, tiles). Round shape images included multiple non-overlapping circles of various sizes and shades ranging from white to black. Rectilinear shape images included diamonds matching the surface areas and shades in the round shape images.

We included three conditions to test animacy and real-world size preferences: animals, big objects, and small objects (145 images each)^22,24^. Each condition included 145 unique categories (e.g., dolphin, raccoon, copier, ladder, bowl, pen). We cropped the object or animal in each image and placed it in the center of the image on uniform gray backgrounds.

To test for mid-level feature preferences, we also created “texform” equivalents of the animals, big objects, and small objects^25,131^. These stimuli are unrecognizable in terms of basic-level category but maintain the second order spatial frequency statistics of the original images^66,67^. We conducted an online behavioral experiment to verify that the texform images obscured recognizability. Participants viewed each image for 1.0s and guessed what they saw. We prompted participants in the following way to encourage responses at a basic-category level: “Be specific with your answers. For example, lion, cup, and bus are all better responses than animal or big object.” At least nine participants categorized each image. We only included images in our image set if at least 75% of the answers provided were unique basic-level categories, ensuring low recognizability of each image. We measured responses to 200 images per category (animals, big objects, and small objects), and after exclusion 145 images per category remained. The previously described image set of recognizable animals, big objects, and small objects corresponded to this culled texform image set.

We included three conditions testing the influence of visual size: faces (100 images), scenes (100 images), and objects (150 images) presented at a smaller visual size. We resized the image content to the middle 1/3 of the width and height of the image, and we made the surrounding image area a uniform gray background.

We included four conditions ranging in spatial scales (100 images each): reachspaces, small scenes, medium scenes, and large scenes. Reachspace images included views of reachable surfaces (e.g., desks, counters, shelves) and were randomly selected from the Reachspace Database^132^. Small-scale scene images included indoor rooms similar in size to bathrooms, bedrooms, and kitchens. Medium-scale scene images included rooms similar in size to art galleries, banquet halls, and lofts. Large-scale scene images included rooms similar in size to arenas, convention centers, and warehouses.

We included four conditions testing sensitivity to spatial layout information (100 images each): empty rooms, rearranged surfaces of rooms, images with single items of furniture, and images with multiple items of furniture^4,19^. We resized images of empty rooms to occupy 70% of the width and height of the full image, then randomly placed them on a black background. For each empty room, a corresponding image was made by rearranging the surfaces into a meaningless configuration. Images with single items of furniture included large objects like cabinets, coffee tables, recliners, and sofas placed on a gray background. Images with four items of furniture included the same items in a 2×2 arrangement on a gray background.

To sample scene conditions with different semantic category and spatial structure^133,134^ we included the following conditions (48 images each): manmade open scenes, manmade closed scenes, natural open scenes, and natural closed scenes.

To measure sensitivity to face shapes across different textural surface properties, we included the following conditions (100 images each): animal faces, cartoon faces, objects which elicit illusory face perception (pareidolia), and matched objects which do not elicit face perception. We cropped animal faces and placed them on a gray background. Animal faces included birds, cats, dogs, fish, horses, pigs, rabbits, and rodents. Cartoon faces were black line drawings in various styles on a gray background. We sampled pareidolia faces from an image set used in previous research^17^. These images include various objects which elicit illusory face perception including buildings, food, and household objects. The last condition included object images which matched the object categories of the pareidolia images, but which do not elicit face perception.

To measure sensitivity to face parts and their arrangement, we created the following conditions (100 images each): typical faces, eyes only, faces without features, faces with rearranged features, and inverted faces^135,136^. We cropped faces and placed them on a gray background. Half appeared to be male, and half appeared to be female. To measure response to the eyes alone, we cropped this region of the face and placed it on a gray background. To create images without facial features, we removed the eyes, mouth, and nose, and layered skin texture over the face. To create faces with rearranged features we placed the eyes, mouth, and noise in a way that disrupted their typical spatial arrangement. Inverted faces were simply the original face images flipped 180 degrees vertically.

### 5.3 Identifying face- and scene-selective model units

We split the localizer image set into two halves: a selection set and a test set. In each ResNet50 model, we measured responses to the selection set at the ReLU outputs of the four residual block groups. For each unit, we conducted two-sample independent t-tests to compare activations between faces vs objects, scenes vs objects, and faces vs scenes. We selected candidate face-selective units which responded more to faces than objects and scenes (p < 0.001 for each contrast). Likewise, we selected candidate scene-selective units which responded more to scenes than faces and objects (p < 0.001 for each contrast).

We evaluated candidate face- and scene-selective units on two test sets from prior research^49,56^. Neural network activations are expressed in arbitrary units and can differ in scale across units, so raw responses are not directly comparable. We therefore z-scored each unit’s responses to place them on a common scale, ensuring equal contribution to mean response estimates. For each image, we measured the mean response across the candidate face-selective units and the mean response across the candidate scene-selective units. Two-sample independent t-tests then compared responses to the target category (faces or scenes) against each of the other categories. For faces, the target categories were adult faces and children’s faces. For scenes, the target categories were scenes, houses, and hallways. Features were classified as face- or scene-selective when responses to the target category exceeded every other category in each t-test.

### 5.4 fMRI Experiment

#### 5.4.1 Participants

Thirty-two healthy adults participated in our experiment (28.6±4.98 years old; mean±sd) All participants had normal or corrected to normal vision. All participants provided written informed consent. Participants received monetary reimbursement at a rate of 12 Euros/hour. Procedures were approved by the ethical committee of the Department of Education and Psychology at Freie Universität.

#### 5.4.2 Experimental Design

Each participant completed two or three runs of a block-design localizer. We presented faces, objects, scenes, and bodies in 16s blocks as participants completed a one-back task. Each run lasted 324s and included five blocks per condition. We asked participants to fixate on a fixation dot throughout the duration of the experiment. Within each block, we presented a new image every second. We presented images for 800ms with an overlaid fixation dot, and then we presented only the fixation dot for 200ms before the appearance of the next image. We presented images at 6.75 degrees of visual angle. Within each participant, we chose a block order which counterbalanced how often each pair of conditions appeared next to each other. Each run included four blocks with just the fixation dot, also lasting 16s each.

Each participant completed ten runs of the main experiment, measuring responses to the test battery. We recorded responses to the 40-condition test battery using a block-design. Participants completed a one-back task and were instructed to maintain fixation on a central fixation dot present throughout the run. Each run lasted 388s and included 20 randomly selected conditions from the test battery. We recorded each condition five times across the full experiment. We presented images in 16s blocks, with 20 images per block. We presented each image for 600ms with an overlaid fixation dot, and then we presented followed only the fixation dot for 200ms before the next image. We displayed images at 6.75 degrees of visual angle. Each run included four blocks with just the fixation dot, also lasting 16s each.

#### 5.4.3 Acquisition Information

We conducted the MRI experiment using a Siemens Magnetom Prisma 3T system (Siemens Medical Solutions, Erlangen, Germany). We collected data with a 64-channel head coil. We measured the anatomical image using a T1-weighted sequence (TR: 1.93s, TE: 3.52ms, number of slices: 256, voxel size: 0.8mm isotropic, FOV: 166.4mm x 237.03mm, flip angle: 8 degrees). We measured functional images with full brain coverage (TR: 2.0s, TE: 30.8ms, number of slices: 72, voxel size: 2.0mm isotropic, matrix size: 110 x 110, FOV: 220mm x 220mm, flip angle: 60 degrees, slice thickness: 2mm, inter-slice gap: 0mm, multiband factor: 3, acquisition order: interleaved).

#### 5.4.4 Preprocessing and GLM Analysis

We preprocessed functional data using fMRIPrep^137^. Data was slice-time corrected to the middle of each TR. A reference volume was generated for head motion correction using a custom methodology of fMRIPrep. Head-motion parameters with respect to the BOLD reference (transformation matrices, and six corresponding rotation and translation parameters) were estimated before any spatiotemporal filtering using mcflirt^138^. The BOLD reference was then co-registered to the T1w reference using mri_coreg (FreeSurfer) followed by flirt^139^ with the boundary-based registration cost-function^140^. Co-registration was configured with six degrees of freedom. Finally, data was smoothed with a Gaussian kernel (FWHM = 4mm).

We estimated single-trial beta weights using GLMSingle^141^. First, we fit a custom hemodynamic response function for each voxel from a predefined library of 20 functions. Second, we conducted data-driven denoising. We identified voxels whose activity did not relate to the experimental design, then we derived a set of noise regressors from their time courses using principal components analysis. We determined the ideal number of noise regressors to include by using cross-validation when fitting the general linear model. Third, we estimated beta values using fractional ridge regression, with cross-validation determining the ideal amount of regularization for each voxel. For each voxel, this procedure yielded a beta estimate for each block across the functional runs.

#### 5.4.5 Defining Regions of Interest

Using data from the localizer runs, we identified the FFA and the PPA in each participant using a Group-Constrained Subject-Specific (GSS) procedure^52^. First, in each voxel in each participant, we conducted two-sided t-tests contrasting beta estimates for the following pairs of conditions: faces vs objects, faces vs scenes, and scenes vs objects. We selected face-selective voxels which responded more to faces than both objects and scenes, using a conjunction threshold of p < 0.0001. Analogously, we selected scene-selective voxels which responded more to scenes than both faces and objects, using the same conjunction threshold. Next, we created group-level parcels by identifying contiguous regions in which at least 10% of participants showed face-selectivity (for the fusiform parcel) or scene-selectivity (for the parahippocampal parcel). Finally, we defined regions of interest in each participant by intersecting each participant’s selectivity map with the corresponding group-level parcel.

### 5.5 Significance Testing for Representational Signatures

Throughout this study, we tested a variety of representational signatures by determining whether the mean activation differed between pairs of conditions in category-selective units (voxels or model units). The structure of our fMRI and neural network data differed, affecting our choices of statistical tests. In the case of the fMRI data, we have a larger number of participants (n = 32) but only five trials per condition per participant, with each trial corresponding to a block of images. This low number of trials was less suitable for statistical testing within each individual, but the higher sample size enabled standard group-level univariate tests typical of fMRI studies with functional regions of interest. In contrast, we have fewer neural network models (n = 8) but many more trials per condition because we measured responses at the image-level in neural network models. This enabled us to conduct statistical tests within each neural network model when comparing activations between conditions.

In the fMRI data, we used group-level binomial tests to compare the mean activation between conditions. For each participant, we computed the mean activation to each condition in each region of interest by averaging beta weights across trials and voxels. To test whether the mean activation differed between pairs of conditions, we performed two-sided binomial tests across participants.

Statistical testing in the neural network models was a two-step process: first conducting within-network statistical tests, then using population prevalence testing to assess whether the effect was common at the group-level. First, we conducted statistical tests within each model for each group of category-selective units (the population of face-selective units or scene-selective units). Akin to measuring the mean response in a region of interest, we computed the mean activation for each image across the group of category-selective units. Then we conducted two-tailed t-tests comparing the image activations between pairs of conditions. This allowed us to determine if each univariate effect was present in each model.

Second, we used population prevalence testing to determine if the group of neural network models exhibited each effect more often than one would expect by chance. This statistical approach is ideal for conducting group-level analyses after within-individual statistics are conducted in small samples^62,142^. In this analysis, we first count the number of networks which exhibited a significant effect. Then we estimated the probability that this number could happen by chance under the null hypothesis that no member of the group exhibited an effect. To estimate this probability, we used the binomial cumulative distribution function (CDF) in the following formula, considering the total number of networks (n), the number of networks exhibiting a significant effect (k), and the probability of making a type I error in each network under the null hypothesis (α):

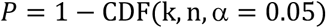

## DATA AND CODE AVAILABLITY

The fMRI dataset collected for this paper is available at https://openneuro.org/datasets/ds007368. The stimulus sets and code for all analyses are available at https://osf.io/d9f5e/overview.

## ACKNOWLEDGEMENTS

This work was supported by a Humboldt Foundation Postdoctoral Research Fellowship (D.J.), German Research Council (DFG) grants (CI 241/1-3, CI 241/1-7, INST 272/297-2) (R.M.C.), and European Research Council (ERC) Consolidator grant (ERC-CoG-2024101123101) (R.M.C.). We thank the Center for Cognitive Neuroscience Berlin for their resources in conducting our fMRI experiment. We thank Johannes Singer and Alessandro Gifford for helpful feedback on the manuscript.

