## Supplementary Material for "Self-supervised learning yields representational signatures of category-selective cortex"

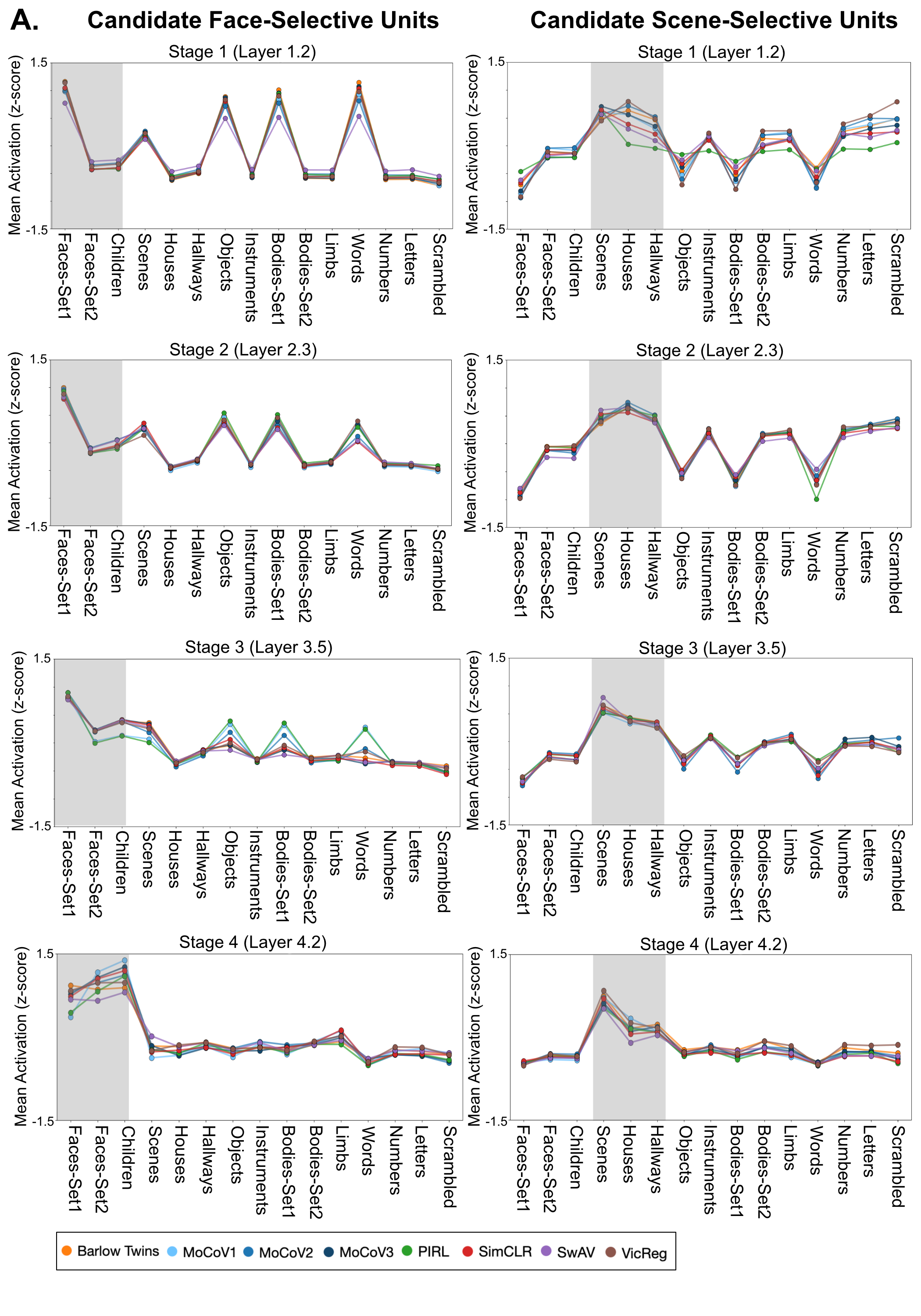

**Figure S1. Responses to the localizer test set in candidate face- and scene-selective units.** A) Candidate face- and scene-selective units were identified in the ReLU output of each stage of each model. Here we depict responses to the test set from early to late layers of the models. Target categories are highlighted in gray (e.g., faces for face-selective units and scenes, houses, and hallways for scene-selective units). T-tests compared the mean activation to these target categories (e.g., faces) to each of the non-target categories (e.g., instruments). For candidate face-selective units (left), models only exhibited full selectivity for face categories in the final model stage (p < 0.01 for each t-test comparing faces to the non-face categories). For candidate scene-selective units, models did not exhibit full selectivity in stages 1 and 2. However, in stages 3 and 4 candidate scene-selective units did respond more to scene categories than all non-scene categories (p < 0.01 for each t-test).

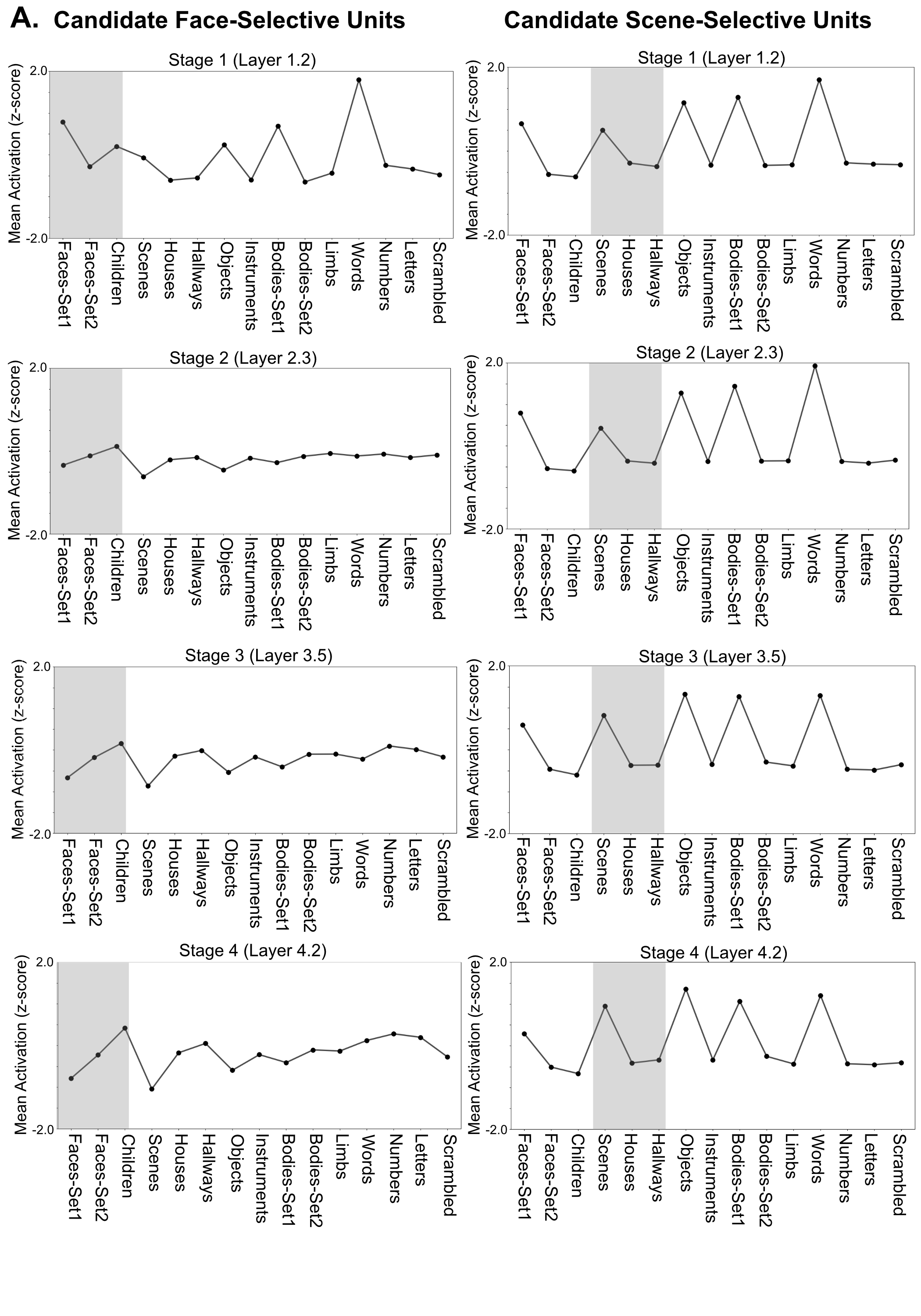

**Figure S2. Responses to the localizer test set in an untrained network.** Responses to the test set are shown for candidate face-selective units (left) and candidate scene-selective units (right) in a ResNet50 network with randomly initialized weights. Images from the target categories are highlighted in gray. Note that the units did not exhibit selective responses for the target categories in any stage of the model.
